# Influenca: a gamified assessment of value-based decision-making for longitudinal studies

**DOI:** 10.1101/2021.04.27.441601

**Authors:** Monja P. Neuser, Franziska Kräutlein, Anne Kühnel, Vanessa Teckentrup, Jennifer Svaldi, Nils B. Kroemer

## Abstract

Reinforcement learning is a core facet of motivation and alterations have been associated with various mental disorders. To build better models of individual learning, repeated measurement of value-based decision-making is crucial. However, the focus on lab-based assessment of reward learning has limited the number of measurements and the test-retest reliability of many decision-related parameters is therefore unknown. Here, we developed an open-source cross-platform application *Influenca* that provides a novel reward learning task complemented by ecological momentary assessment (EMA) for repeated assessment over weeks. In this task, players have to identify the most effective medication by selecting the best option after integrating offered points with changing probabilities (according to random Gaussian walks). Participants can complete up to 31 levels with 150 trials each. To encourage replay on their preferred device, in-game screens provide feedback on the progress. Using an initial validation sample of 127 players (2904 runs), we found that reinforcement learning parameters such as the learning rate and reward sensitivity show low to medium intra-class correlations (ICC: 0.22-0.52), indicating substantial within- and between-subject variance. Notably, state items showed comparable ICCs as reinforcement learning parameters. To conclude, our innovative and openly customizable app framework provides a gamified task that optimizes repeated assessments of reward learning to better quantify intra- and inter-individual differences in value-based decision-making over time.

## Introduction

Learning from past experiences is essential to optimize decision-making and adaptive behavior. Reinforcement learning models provide useful quantifications of individual choice behavior and the integration of information over repeated decisions (Sutton & Barto, 2018). Hence, disturbances in reward learning may result in maladaptive choices and are associated with various mental and metabolic disorders, such as depression (Chen, Takahashi, Nakagawa, Inoue, & Kusumi, 2015; Eshel & Roiser, 2010; Mkrtchian, Aylward, Dayan, Roiser, & Robinson, 2017), eating disorders (Schaefer & Steinglass, 2021), and obesity (Coppin, Nolan-Poupart, Jones-Gotman, & Small, 2014; Kroemer & Small, 2016). Parameters of individual reinforcement learning, such as the learning rate or reward sensitivity may even serve as transdiagnostic biomarkers (Montague, Dolan, Friston, & Dayan, 2012) for aberrant cognitive processes that contribute to key symptoms of disorders, such as apathy or anhedonia (Husain & Roiser, 2018; Huys, Pizzagalli, Bogdan, & Dayan, 2013). In the light of growing interest in reinforcement learning for psychological diagnostics, it is worth noting that the effective use of measures as biomarkers for prediction and classification of mental function requires a thorough evaluation of their psychometric properties (Fröhner, Teckentrup, Smolka, & Kroemer, 2019; Hedge, Powell, & Sumner, 2018; Moriarity & Alloy, in press). However, since most studies are conducted in laboratory settings with limited numbers of participants and repeated measures, a systematic evaluation of the psychometric properties of reinforcement learning parameters, such as their test-retest reliability, is still lacking.

To overcome practical limitations of scale of lab-based testing, online and smartphone-based assessments are becoming increasingly popular. Once a setup is established and verified, they enable data acquisition of many individuals across multiple time points embedded in different situations (Gillan & Rutledge, 2021). These aspects facilitate longitudinal data collection and may help improve generalization to robust behavioral predictions outside of the laboratory. Especially in the domain of mental health, methods such as ecological momentary assessment (EMA) or experience sampling are increasingly common to monitor fluctuations in mood or other psychological and physiological states (e.g., Blain & Rutledge, 2020; Ebner-Priemer & Trull, 2009; Killingsworth & Gilbert, 2010; Mason et al., 2018; Perrez, Schoebi, & Wilhelm, 2000; Wonderlich et al., 2018). Consequently, assessing such fluctuations in mental states over time may help predict the onset of disorder-related behavior which is impossible to recreate in the lab, such as binge eating (Smyth et al., 2009; Svaldi, Werle, Naumann, Eichler, & Berking, 2019; Wonderlich et al., 2018) or binge drinking (Dvorak & Simons, 2014; Shiffman, 2009; Wray, Merrill, & Monti, 2014). Beyond practical aspects of data collection, smartphone-based assessments may also reach a more diverse population of users, which increases the variance between participants and improve generalizability (Gillan & Rutledge, 2021; Spook, Paulussen, Kok, & Van Empelen, 2013). Such an online format has been used for reinforcement learning tasks as well, leading to big samples that contributed to revisions of commonly used models (Gillan, Kosinski, Whelan, Phelps, & Daw, 2016; Rutledge, Skandali, Dayan, & Dolan, 2014). However, a similar innovation with repeated assessments has yet to materialize. To conclude, online-based assessments of reinforcement learning may provide a powerful means to collect ecologically and psychometrically valid estimates to predict individual trajectories, but this requires a framework for the repeated collection of decision-related parameters over time.

To interpret individual trajectories, including potential effects of interventions or as markers of clinical progression (Dubois & Adolphs, 2016), a sufficiently high level of test-retest reliability is necessary. Reliability is a prerequisite of a paradigm’s validity (Cronbach & Meehl, 1955) and a lack of the formal assessment of reliability might hamper the widespread application of reinforcement learning tasks in the study of psychopathology (Rodebaugh et al., 2016). Amongst common decision-making tasks, reliabilities range from *r* = .15 to .69 (Buelow & Barnhart, 2018). For common probabilistic reinforcement learning tasks, parameter estimates of computational models, such as the learning rate, only show low to moderate reliability. For example, Santesso et al. (2008) reported a test–retest correlation for reward learning of *r* = .50 in a probabilistic reward learning task. In a go-no-go reinforcement learning task, Moutoussis et al. (2018) reported low to moderate test-retest correlations (range *ρ*: .07 [noise parameter] to .37 [model fit]) of parameter estimates. In clinical samples, even lower reliabilities (mean ICC = .33, range: .08 -.58) have been reported for such tasks, for example, in individuals suffering from schizophrenia (Pratt et al., 2020; Shiner et al., 2012). Based on online assessments of probabilistic reinforcement learning and reversal learning tasks, Weidinger, Gradassi, Mollemann, and van den Bos (2019) reported fair to good ICCs for basic outcome measures (i.e., task accuracy or win-stay: ICC between .43 - .76) and learning rate estimates (ICC = .54) (Weidinger et al., 2019). Likewise, measures of self-regulation show poor to moderate reliability in online behavioral assessments (median ICC = 0.311, IQR = −0.091 - 0.665), which is lower compared to questionnaire surveys (mean ICC = 0.716) and published ICCs for lab-based assessment (mean ICC = 0.610 (Enkavi et al., 2019)) that could be, however, overly optimistic. Notably, raw dependent variables (i.e., response time and accuracy) had comparable reliabilities to latent variables (i.e., model estimates such as drift rate; (Enkavi et al., 2019)). Taken together, compared to questionnaire measures and despite the widespread use of reward learning tasks, there is insufficient information on the reliability of potential biomarkers related to value-based decision-making and reward learning, and the few studies point to low or, at best, fair test-retest reliability.

To summarize, cross-sectional measures provide a snapshot of value-based decision-making, which might be of limited use for future applications to predict individual trajectories, specifically if they lack sufficient test-retest reliability. Consequently, psychometric evaluations of repeated assessments of alleged biomarkers are crucial to uncover the psychobiological processes contributing to key symptoms of mental disorders. Here, we present an innovative cross-platform open-source app (Influenca, www.neuromadlab.com/en/influenca-2), designed to assess value-based decision-making and reward learning over weeks on the participants’ preferred devices. By measuring decision-making over extended time periods, we can also evaluate the reliability of behavioral parameters with much higher precision. Better knowledge about the parameters’ psychometric characteristics may improve future clinical decisions (e.g., targeted prevention or treatment monitoring) regarding mental disorders that are characterized by dysfunctions in value-based decision-making.

## Methods

### Participants

The initial sample included 294 individuals who downloaded and played the app until 15^th^ of October 2020. For the current analyses, we included all participants that completed at least 10 runs of Influenca surpassing our data quality screening (i.e., 316 runs were excluded due to random or deterministic behavior; 5 < log-likelihood < 102.58). This led to a final sample of N = 127 participants (*M*_age_ = 35.78 years, *SD* ± 14.13) with 2904 valid runs.

The collected data was part of a study approved by the local ethics committee and was conducted in accordance with the ethical code of the World Medical Association (Declaration of Helsinki). Informed consent was obtained twice, depending on the stages of the study. First, all participants provided informed consent by clicking a checkbox (Kraut et al., 2004) when registering for the app, stating they agree with the terms of service and usage of pseudonymized and anonymized data for the specified scientific objectives. Second, participants were provided with a second informed consent form before completing the online questionnaires. Participants who completed the app as part of a clinical study (n = 99) received a fixed compensation (€20) for the complete online assessment, which included questionnaires.

### Influenca and reinforcement learning game

To repeatedly assess reinforcement learning, we developed the cross-platform app Influenca. The app includes 31 runs of a reinforcement learning game based on a classic paradigm (Behrens, Woolrich, Walton, & Rushworth, 2007) with changing reward probabilities. Before each run, participants completed several EMA items capturing momentary metabolic (hunger, satiety, thirst, time since last meal, consumption of coffee or snacks in the last two hours) and mental states (alertness, happiness, sadness, stress, distraction by environment, distraction by thoughts). Responses were given using either visual analog scales (VAS: hunger, fullness, thirst, alertness, happiness, sadness, stress, distraction by environment, distraction by thoughts), Likert scales (last meal), or binary scales (snack, coffee, binges).

In the gamified scenario of Influenca, participants had to fight a virus pandemic by finding the most effective medication. In each run, participants were presented a new virus. In each trial, they had to choose between two medications depicted as syringes of different colors with (initially) unknown win probabilities (Figure 1). Notepads corresponding to each syringe showed the number of people cured in the trial, if the chosen medication was correct (“win”). The corresponding points were added to the total score. If the chosen option resulted in a loss, the points were subtracted from the total score instead. A counter in the upper left corner showed the number of completed trials and the current total score.

**Figure 1:**
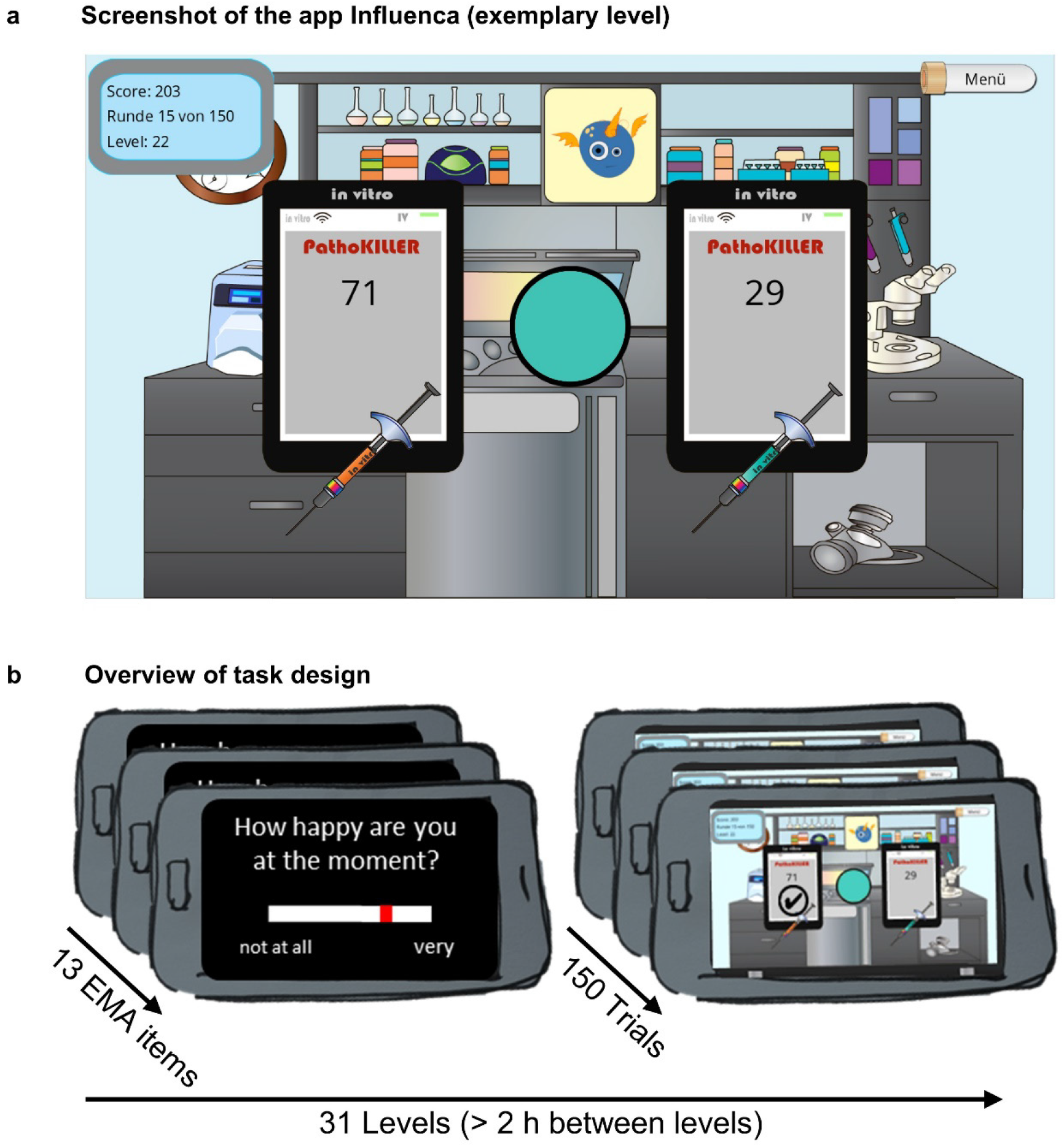
Illustration of the reinforcement learning task design. **A**. Representative in-game screen of Influenca. To earn points, participants must identify which medication is most effective in fighting pathogens. In each trial, only one drug is effective to cure people. In this trial, the orange drug could treat 71 people and the turquoise drug would only treat 29 people. The circle depicts the color of the drug that was effective in the previous trial. If participants pick the correct medication, their score increases by the number of cured people (win). In contrast, if they pick the incorrect medication, the number of falsely treated people will be subtracted from the score (loss). **B**. Procedure of the Influenca levels. Each level starts with EMA questions about participants’ current mood and other states prior to the actual game, followed by 150 trials of the reinforcement learning paradigm. To ensure sampling across different states, there was a minimum of 2 hours waiting time enforced between the runs.

In each trial, the number of points added up to 100 across both options. Win probabilities were independent of reward and added up to 1 across both options. Since participants repeated the task up to 31 times, win probabilities of the options were determined by a Gaussian random walk algorithm and thus fluctuated over time. The use of random walks was intended to reduce meta-learning about when reversals or changes in contingencies would occur (Boehme et al., 2017; Hämmerer et al., 2019). Each run was randomly initialized with a “good” (*p*_*win*_ = 0.8) and a “bad” (*p*_win_ = 0.2) option. To encourage replay, participants had to fight against a new virus in each run. After completing a run, the defeated virus was added to a scoreboard and each level highlighted the scientists’ increased “prestige” by showing an improved quality of the lab equipment, as depicted in the game’s graphics.

### Experimental procedure

Participants installed the app on their preferred device by obtaining the installer file from our homepage (https://neuromadlab.com/en/influenca-2/). They provided a mail address at registration for app-specific communication (e.g., sending automated reminder mails, to send the individualized link for the questionnaires and the activation code to unlock level 11 to 31 after completing the online questionnaires). To ensure confidentiality, the mail address was stored apart from the experimental data.

Before starting the first run, participants were asked to read through a detailed instruction explaining the controls as well as the game’s cover story and rationale. They could re-read this instruction at any time by opening it via the game’s menu. A version of this instruction was also posted at the download section on our lab homepage. Participants were instructed to play at different times throughout the day and in different (metabolic) states to sample data in diverse situations to improve generalizability. To ensure sufficient distinctiveness across runs, we required a delay between runs of at least 2 h, but there was no time restriction to complete the 31 runs. Collected data was stored locally on the participant’s device and, once connected to the Internet, synchronized with a database located at the Department of Psychiatry and Psychotherapy, University of Tübingen.

### Data analysis

#### Reinforcement learning model

To model different facets of reward learning, we used choice data from individual runs and estimated individual learning parameters: learning rate (α), reward sensitivity (β), and risk aversiveness (γ) using an established reinforcement learning model (Behrens et al., 2007; Neuser, Kühnel, Svaldi, & Kroemer, 2020). In this model, participants are assumed to decide between the options in each trial based on the inferred win probability of each option. These probability estimates, *p*_*win*_, are learned, by integrating over outcomes of previous decisions following a simple delta rule:

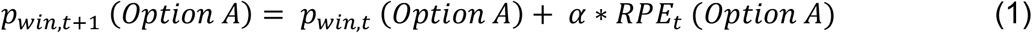

where α ∈ [0, 1] denotes the learning rate that quantifies the speed of adaptation of an individual’s choice preference along with changing outcome contingencies. In other words, high learning rates lead to quick updates by putting more weight on recent choice outcomes. In contrast, low learning rates lead to slow updates by putting less weight on recent choice outcomes and more weight on former feedback (Figure 2a). Formally, the learning rate scales the reward prediction error, RPE, that describes the difference between the choice outcome, *r*, and the expected probability of winning for a chosen option:

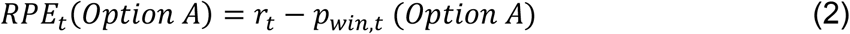

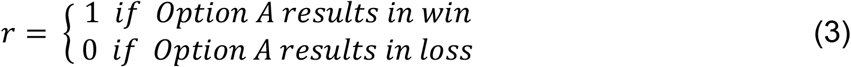

**Figure 2.**
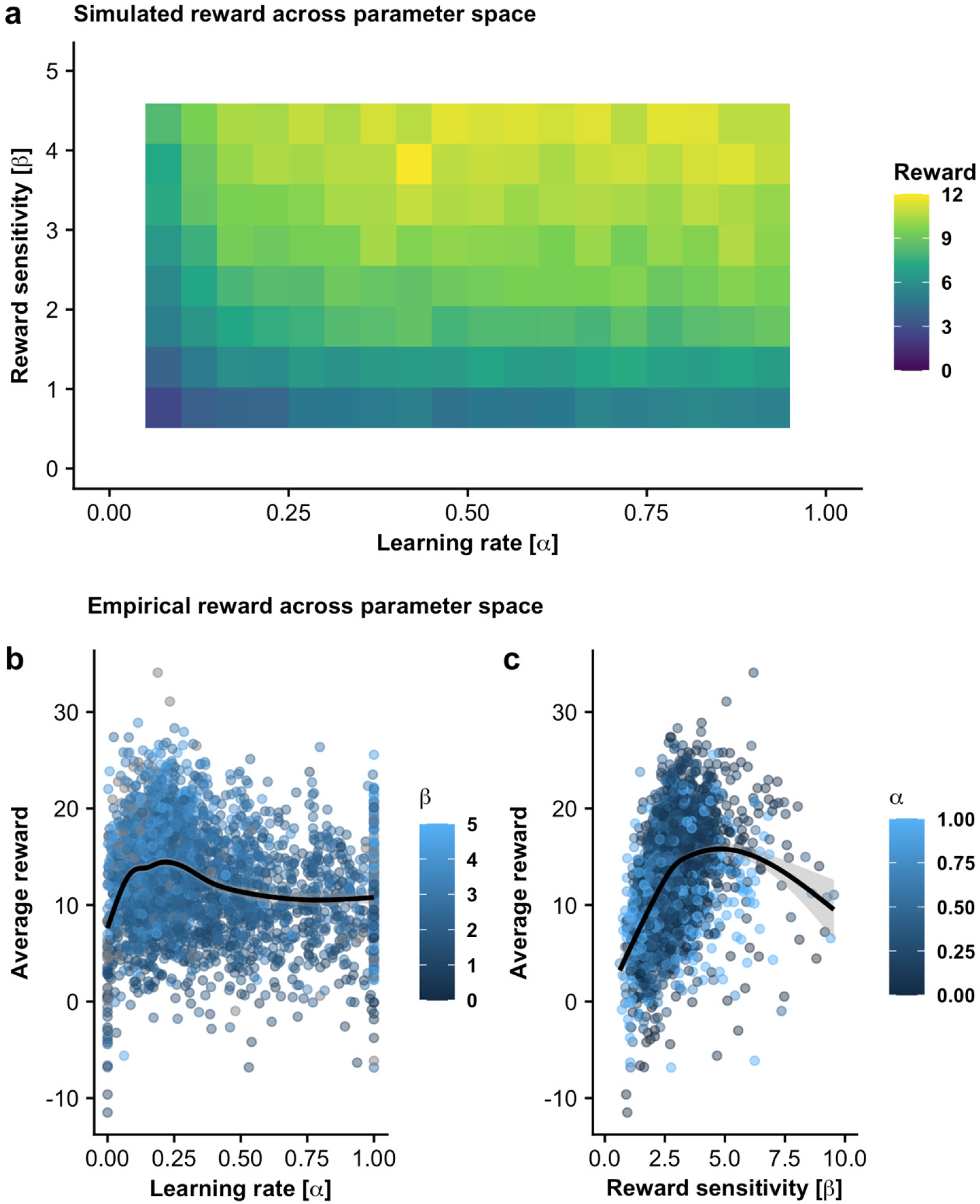
Reward outcome per run for different combinations of learning parameters based on simulated and behavioral data. **A**. Simulation of N = 50,000 players shows high rewards for different combinations of learning rate and reward sensitivity. **B**. Empirical data from the participants show high average rewards for moderate learning rates and moderate reward sensitivity.

Choices in each trial are then generated based on the estimated win probability of the option (*p*_*win,t* +1_ (*Option A*)) and the associated reward values of each option (*f*) (*f*_*A*_ ∈ [0,100] and *f*_*B*_ ∈ [0, 100] *with f*_*A*_ = 100 − *f*_*B*_). In the optimal case, an individual chooses the option that maximizes the expected outcome by scaling the win probability of each option with its associated reward into action weights (*W*)

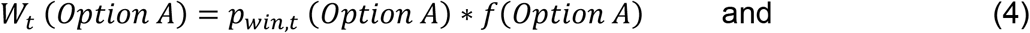

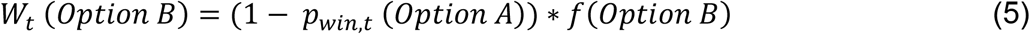

However, it is unlikely that all individuals always act in a rational manner, as some may be more risk prone and decide predominantly based on the win probability while ignoring the associated rewards at stake. In contrast, other individuals may choose the potentially more rewarding option even if it is less probable to lead to a win. Differences in this evaluation can be quantified by an additional parameter, γ, representing the individual risk aversiveness:

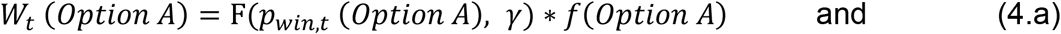

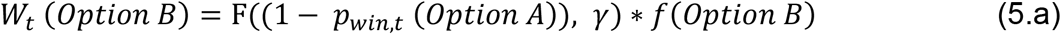

Here, *F* is a linear transform that scales the win probability according to the individual level of risk aversiveness yet ensures that it remains within the bounds of 0 and 1.

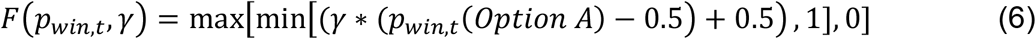

The previously described case of rational decision-making would be achieved with γ = 1, whereas γ > 1 would lead to risk-averse behavior since decisions are predominantly based on the learned probabilities and less on differences in reward points. In contrast, γ < 1 would make differences in reward points more important. Finally, the probability to choose a given option is computed with a sigmoidal probability function based on the action weights.

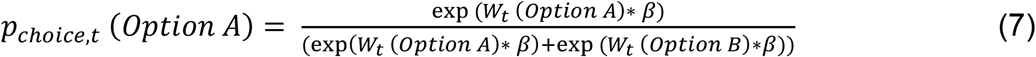

Note that this function contains the reward sensitivity, *β* ∈ [0, Inf], that scales the subjective value. The reward sensitivity indicates the predictability of value-based decisions in such a way that high reward sensitivity leads to very predictable choices based on the action weights, while low reward sensitivity leads to noisier decisions (Neuser et al., 2020).

We fit the model using maximum likelihood estimation with the *fmincon* algorithm implemented in MATLAB 2018b. We previously showed that parameters could be successfully estimated in simulated data (Neuser et al., 2020) in case of rational decision making (γ = 1).

Since many reinforcement learning models only include learning rate and reward sensitivity parameters, we compared this basic reinforcement model with fixed risk aversiveness (γ = 1; assuming rational integration of win probability and expected reward) with a model including γ as free parameter. The likelihood ratio test (*lratiotest;* LR) based on the log-likelihood of the model fit provided strong evidence for the extended model including risk aversiveness (*LR* = 43624, *p* < .001, *df* = 2904), confirming a better fit for 108 out of 127 participants. Thus, we performed all following analyses using the extended model including all three parameters.

To ensure that the choice behavior of participants was sufficiently well approximated by the model, we used the log-likelihood of each run as criterion. We excluded 316 runs (8.3 % of all runs) with a poor model fit due to random or invariant choices (e.g., always option A) from further analyses (5 < log-likelihood< 102.58, boundaries estimated from simulated data with random choices). As model-independent performance measures, we used the average of earned points per trial for each run and response times for each decision. Trials with extreme response times (50 ms < response time < 10,000 ms) were excluded from the response time analysis (5159 trials, 0.9% of all trials).

#### Run effects

To estimate effects of repeatedly playing the game on estimated parameters, we used linear mixed-effects models (lmerTest; Kuznetsova, Brockhoff, & Christensen, 2017). We predicted the estimated behavioral parameters and model fit using the log-transformed run number as fixed effect. To account for inter-individual differences, run number and the intercept were modeled as random effects.

#### Test-retest reliability

To assess the reliability of behavioral parameters and state items, we estimated ICCs using linear mixed-effects models (Raudenbush & Bryk, 2002). The ICC describes the reliability of a measure on the scale of a correlation coefficient, where values close to 1 reflect high similarity within participants (“classes”, denoted as ID), whereas lower ICCs indicate lower similarity within participants. As described in Raudenbush and Bryk (2002), we derived unconditional ICC based on the null model: Parameter ∼ (1|ID) with the formula:

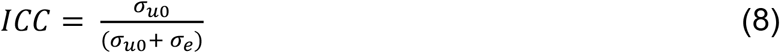

where *σ*_*uo*_ is the variance explained the random intercept (ID) and *σ*_*e*_ denotes the residual variance. Additionally, we calculated the conditional ICC taking (fixed) run effects (log-transformed) into account with the following mixed-effects model:

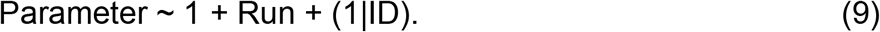

We interpreted ICC values according to recommendations by Shrout and Fleiss (1979) so that values < 0.4 reflect poor, values between 0.4 and 0.6 reflect fair, between 0.6 to 0.75 reflect good, and values > 0.75 reflect excellent reliability. Lastly, to better describe which runs within the game provide the most reliable parameter estimates, we calculated the test-retest rank correlation (Spearman) for each run with the average of the respective parameter across all other runs. We interpreted the correlations according to recommendations by Taylor (1990) for correlation coefficients. Here, *r*s < .35 reflect low, *r*s between .36 and .67 reflect modest, *r*s > .67 reflect high correlations.

#### Statistical threshold and software

All statistical tests were performed using a significance level of α = 0.05 (two-tailed). Data preprocessing was done with MATLAB 2020a. Linear mixed-effects models and ICCs were estimated in R Studio (R Version 3.5.3; R Core Team, 2014). Plots were created in R (R Version 3.5.3; R Core Team, 2014) using the package ggplot2 (Wickham, 2011).

## Results

### Validation of parameter estimates and game mechanics

In each run, participants made 150 choices between two options of varying reward value and fluctuating win probabilities that had to be inferred over time. To estimate reinforcement learning parameters from value-based choices, we applied computational modeling. The task was designed to encourage expression of inter-individual differences in reward learning by providing a large plateau of moderate to high average rewards per run across a wide parameter space (Figure 2a). Comparable patterns of average reward across different combinations of estimates are also evident in empirical data (Figure 2b and 2c), albeit with a narrower range compared to simulations that do no entail correlations of estimated parameters (Daw, 2011). We see an increased average reward for moderate learning rates α ∼ 0.20 (Figure 2b), indicating that integrating over multiple decisions is advantageous to track “true” fluctuations in the hidden random walks. This basic insight is illustrated in Figure 3a, where a high learning rate (top panel) leads to fast switches in choices due to recent events. In contrast, a lower learning rate (Figure 3a, bottom panel), leads to more patience in light of surprising choice outcomes reducing the number of sharp transitions between preferred options.

**Figure 3.**
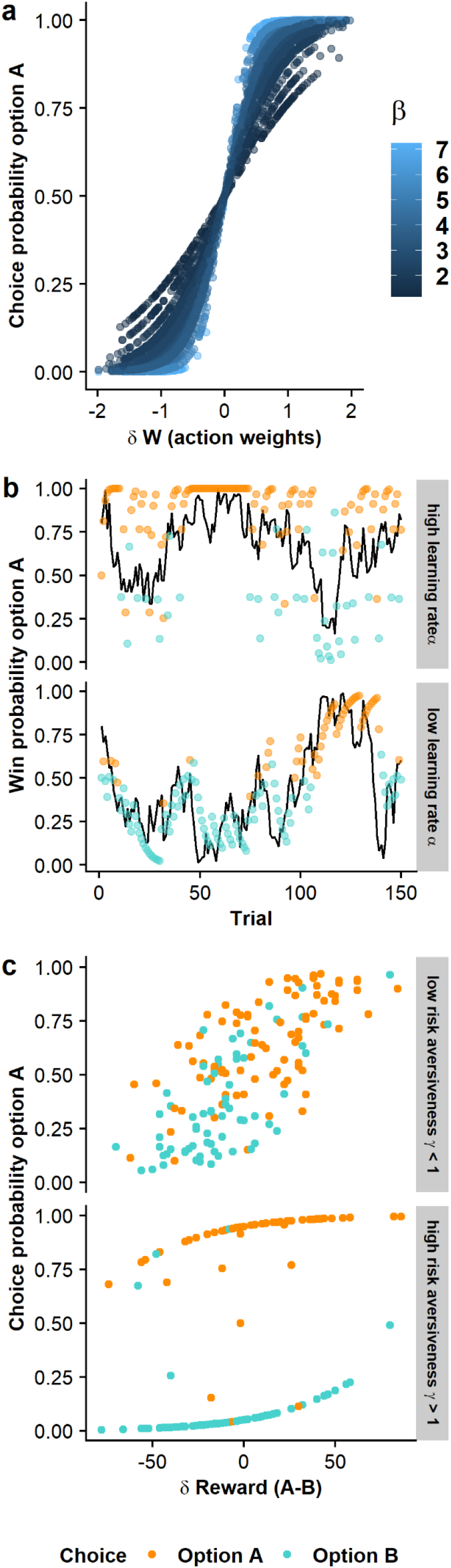
Illustration of estimated parameters and their relation to differences in value-based decision-making and learning (representative participants). **A**. The reward sensitivity beta scales how action weights (i.e., a combination of estimated probability and potential reward value) are translated into choices. Higher reward sensitivities translate to more deterministic choices (i.e., exploitation), whereas lower reward sensitivities lead to more random choices (i.e., exploration). **B**. The learning rate alpha captures how quickly estimated win probabilities are updated in light of new information. High learning rates (upper panel) lead to fast updates and quick forgetting of long-term outcomes. The black line depicts the latent win probability, while the points depict the estimated win probability based on the reinforcement learning model. **C**. Risk aversiveness scales the importance of the estimated win probability of each option compared to the offered rewards. Low values (<1, upper panel) reduce the importance of the learned win probabilities leading to choices based primarily on the potential reward at stake. In contrast, high values (>1, lower panel) increase the importance of the learned win probabilities. If the parameter is very high, it may lead to very deterministic choices, as small differences in the inferred probability to win are exaggerated.

In line with simulated results, our data shows a clear association of higher average rewards with higher reward sensitivities, although the improvement plateaus around the upper limit of our simulation (β > 4.5). Since reward sensitivity reflects the predictability of choices based on inferred differences in values, a high value leads to more deterministic choices even for small relative differences in the values of the options. In contrast, reward sensitivity close to zero corresponds to random decisions.

The risk aversiveness parameter reflects whether participants put more emphasis on the win probability than the value. A low gamma parameter corresponds to low risk aversiveness (Figure 3c, upper panel), indicating that choices are mainly driven by the potential value of the outcome. A high gamma parameter corresponds to high risk aversiveness (Figure 3c, lower panel), indicating that choices are mainly driven by the inferred win probability. Notably, almost all participants showed a higher than rational risk aversiveness (96%) suggesting they weighed option values less and focused more strongly on the probability to win than optimal when making a choice.

Conditional and unconditional ICCs of average rewards per run were lower compared to the parameter estimates (unconditional and conditional ICC = 0.09), indicating little grouping of data within individuals compared to overall variability across runs. Response times ranged from 0.054 to 9.9 s with higher unconditional and conditional ICCs compared to average reward rates (0.33 and 0.38, respectively; Table 1).

**Table 1.** Descriptive statistics and reliabilities of dependent variables

| Measures | Mean | SD | Median | 10<sup>th</sup><br>Percentile | 90<sup>th</sup><br>Percentile | ICC<sub>unc</sub> | ICC<sub>cond</sub> |
| --- | --- | --- | --- | --- | --- | --- | --- |
| **Behavioral indices** |  |  |  |  |  |  |  |
| Response time [s] | 0.99 | 0.69 | 0.85 | 0.52 | 1.79 | 0.33 | 0.38 |
| Wins | 12.63 | 5.55 | 12.72 | 5.66 | 19.58 | 0.09 | 0.09 |
| **Model parameters** |  |  |  |  |  |  |  |
| Log-likelihood | -52.93 | 19.10 | -53.05 | -78 | -28.7 | 0.34 | 0.31 |
| Learning rate | 0.36 | 0.27 | 0.26 | 0.08 | 0.83 | 0.49 | 0.52 |
| Reward sensitivity | 2.87 | 1.16 | 2.70 | 1.67 | 4.17 | 0.36 | 0.37 |
| Risk aversiveness | 5.45 | 6.28 | 2.71 | 1.11 | 14.05 | 0.22 | 0.22 |
| **State items** |  |  |  |  |  |  |  |
| Alertness | 56.99 | 23.00 | 58 | 26 | 88 | 0.29 |  |
| Happiness | 59.82 | 24.03 | 62 | 26 | 91 | 0.45 |  |
| Sadness | 27.84 | 24.69 | 21 | 1 | 67 | 0.43 |  |
| Stress | 29.42 | 24.21 | 24 | 1 | 65 | 0.47 |  |
| Distraction by environment | 26.10 | 23.19 | 20 | 1 | 61 | 0.39 |  |
| Distraction by thoughts | 30.55 | 25.06 | 24 | 2 | 68 | 0.53 |  |

Note: SD = standard deviation, ICC = intraclass correlation coefficient

### Reinforcement learning improves over runs

To investigate changes in reinforcement learning over runs, we estimated a mixed-effects model for each behavioral parameter, including intercept and log(run) (random intercept and slopes). Repeatedly playing the game led to decreased reaction times (*b* = -314 ms, *t* = 20.03, *p* < .001) indicating increased familiarity with the task. In general, participants successfully learned the correct choice within the task as indicated by a positive average reward obtained (first run: *M* = 8.98, *SD* = 5.85; all runs: *M* = 12.63, *SD* = 5.55), and these rewards increased over runs (*b* = 1.37, *t* = 9.53, *p* < .001). Moreover, the decrease in response times and increase in reward were negatively correlated (*r* = -.25; *p* < .001) suggesting that increased proficiency also speeds up decisions without a detrimental effect on accuracy. The improvement in model-independent performance indices was mirrored in the parameter estimates. Over runs, the learning rate decreased (Figure 4a, *b* = -0.08, *t* = -9.00, *p* < .001) and the reward sensitivity increased (Figure 4b, *b* = 0.24, *t* = 3.95, *p* < .001), both associated with higher average reward (Figure 2a-b). Moreover, the model fit (log-likelihood) also increased over runs (Figure 4d, *b* = 6.87, *t* = 11.09, *p* < .001), suggesting that choices became more aligned with the estimated reinforcement learning model. Nonetheless, participants became more risk averse over runs (Figure 4c; *b* = 0.73, *t* = 3.55, *p* < .001), even if this led to greater deviations from rational behavior.

**Figure 4.**
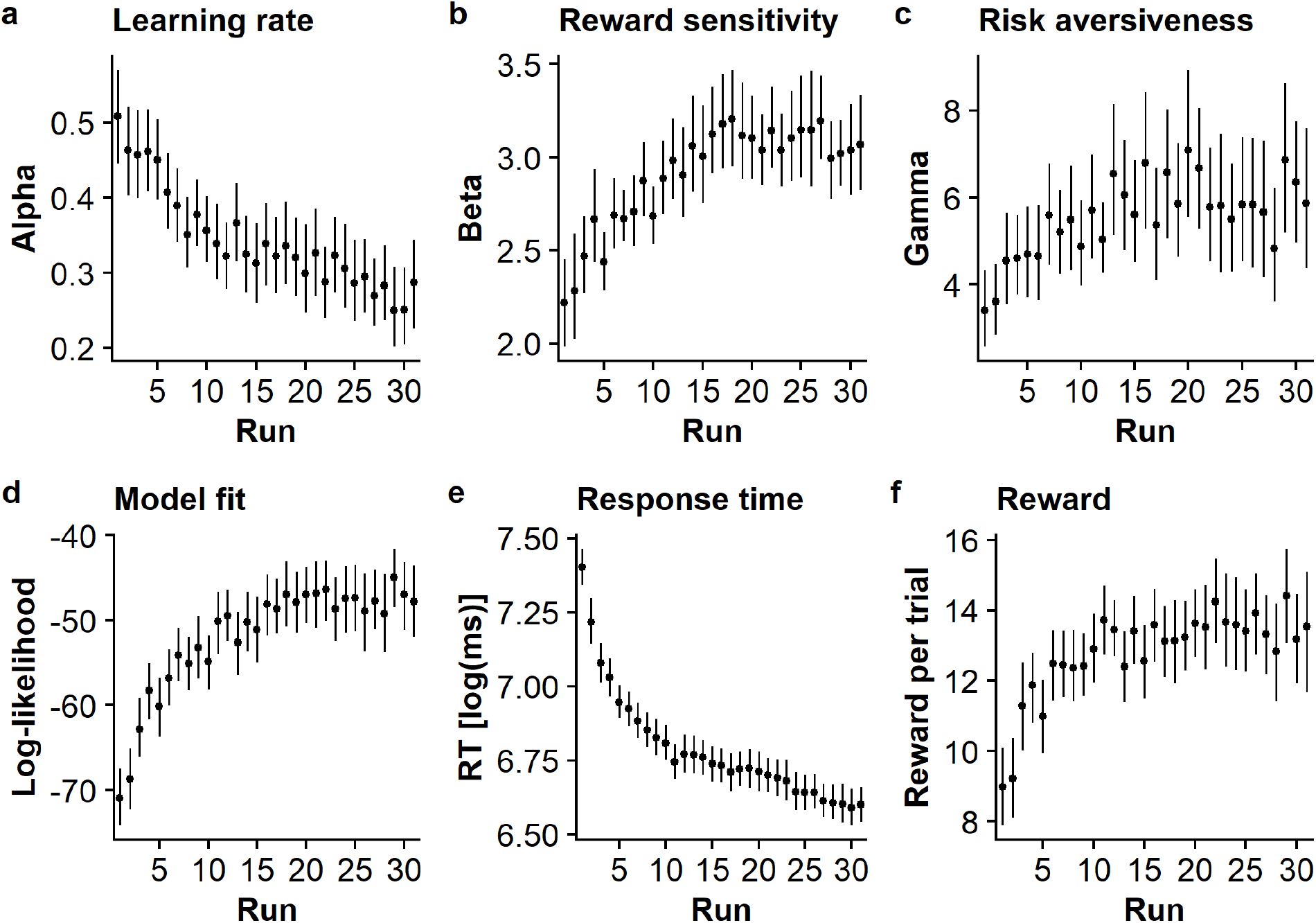
Reinforcement learning parameters and model-independent performance indices improve over runs. **A**. Learning rate, α, decreases over runs (*b* = -0.08, *p* < .001). **B**. Reward sensitivity β increases over runs (*b* = 0.24, *p* < .001). **C**. Risk aversiveness, γ, increases over runs (*b* = 0.73, *p* < .001). **D**. The model fit (log-likelihood) improves over runs (*b* = 6.87, *p* < .001). **E**. Response times decrease over runs (*b* = -314 ms, *p* < .001) and **F**. The average reward increases over runs (*b* = 1.37, *p* < .001). The dots show mean values with 95% bootstrapped confidence intervals.

### Reliability of reinforcement learning parameters is comparable to momentary states

To assess the reliability of reinforcement learning parameters, we computed unconditional and conditional ICCs adjusting for the behavioral adaptation over runs (Figure 4), yielding fair ICCs for learning rates (unconditional ICC = 0.49, conditional ICC = 0.52) and poor ICC values for both reward sensitivity (unconditional ICC = 0.36, conditional ICC = 0.37) and risk aversiveness (unconditional and conditional ICC = 0.22; Table 1). To investigate whether early or late runs are more reliable, we calculated the Spearman rank correlation for each run with the average of all other runs. Notably, test-retest correlations of early runs were lower compared to late runs and improved until reaching a plateau after approximately 7 runs (Figure 5). This indicates that behavior within the task becomes more reliable with higher task proficiency, suggesting that late runs are better estimates of the “typical” performance on the task compared to early runs.

**Figure 5.**
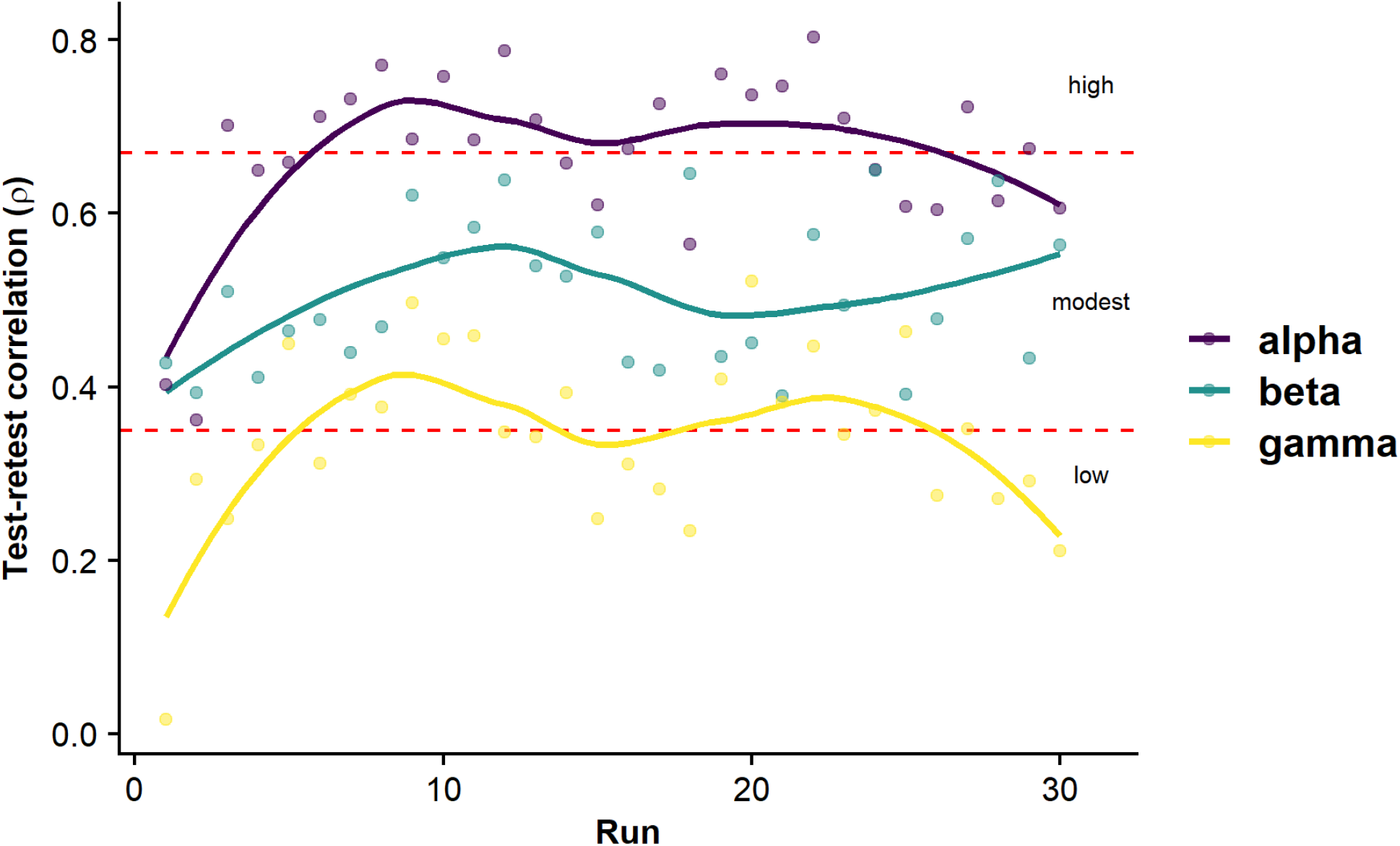
Reliability of parameter estimates from the learning model improves after the first runs. Dots depict the correlation of parameter estimates in one run with the mean across all other runs (leave-run out), separated per parameter. Red dashed lines show the classification of correlation magnitudes according to Taylor (1990).

To relate the reliability of reinforcement learning parameters with the reliability of other measures (i.e., momentary states), we analyzed a subset of state items that are assessed prior to each run. We calculated descriptive statistics and ICCs of EMA items alertness, happiness, sadness, stress, distraction by environment, and distraction by thoughts (Table 1, Figure 6). The ICCs of the EMA items ranged between poor (alertness: 0.29, distraction by environment: 0.39) and fair (sadness: 0.43, happiness: 0.45, stress: 0.47, distraction by thoughts: 0.53) values. Of note, they were in a comparable range as the reinforcement learning parameters. Illustratively, items reflecting emotional states (happiness, sadness, stress) had similar ICCs as the learning rate, the most reliable reinforcement learning parameter. These results point to a comparable ratio of within- and between-subject variance for behavioral parameters as for alleged state items.

**Figure 6.**
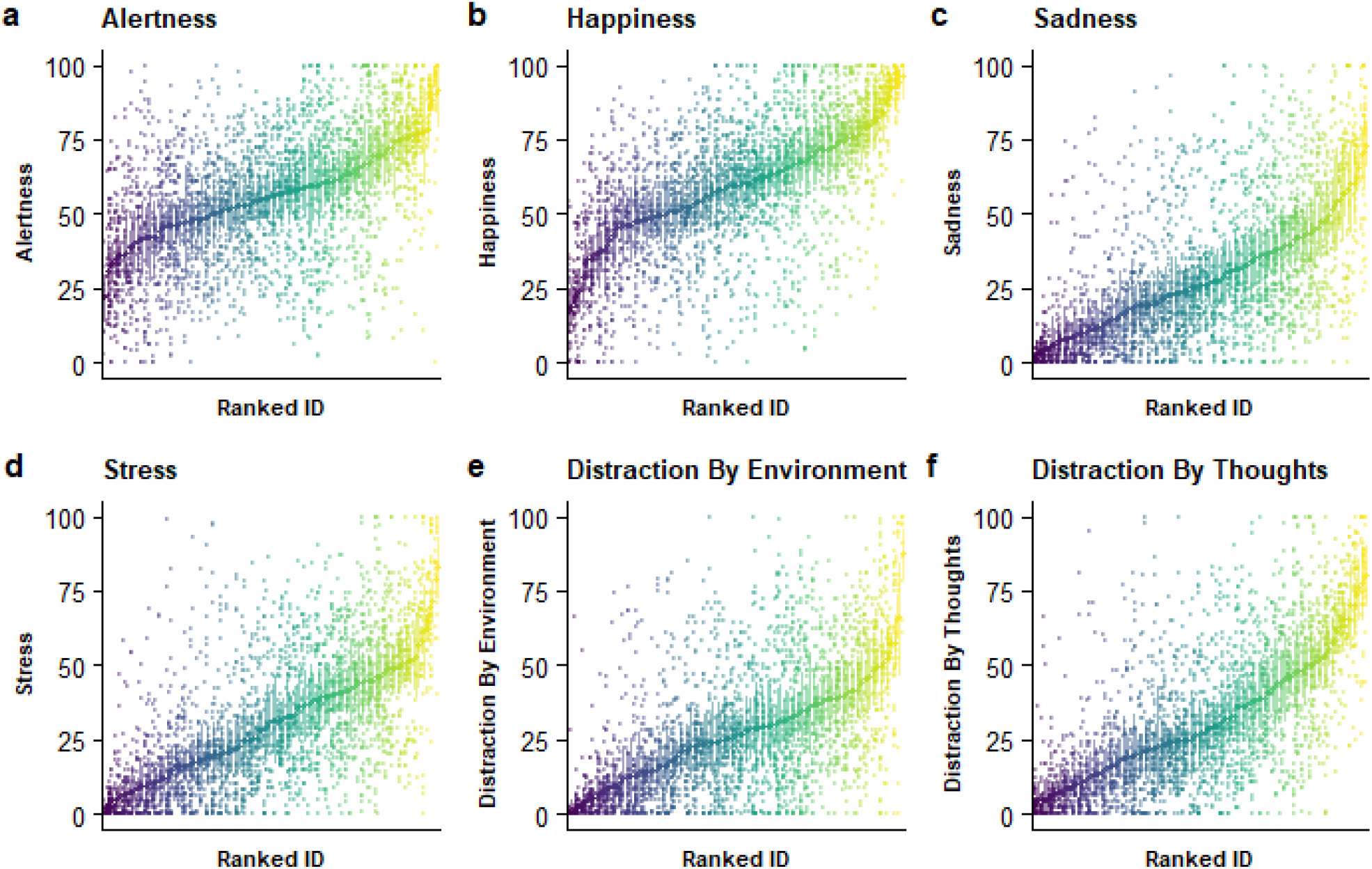
State items show poor to fair intra-class correlation coefficients (0.29-0.53). **A-F**. Mean and variance of selected EMA items per individual, ranked by mean value of each participant across runs. Dots represent values per run as entered via VAS.

## Discussion

Value-based decision-making and learning are integral parts of adaptive behavior. Since alterations have been linked to multiple mental disorders, it is pivotal to develop accessible tools that capture reliable inter-individual differences in value-based decisions to propel the use of deeply phenotyped behavioral data for clinical classification and prediction. To this end, we introduce our open-source cross-platform application Influenca, which comprises multiple runs of a reward learning task and (customizable) EMA state items. Using preliminary data of 2904 runs from users with a minimum of ten runs, we show that our app provides detailed insight into the reliability of common indices of value-based decision-making and learning over extended periods of time. In line with the few available reports of test-retest reliability in comparable lab-based assessments, the reliabilities of the estimated model parameters were poor to fair, suggesting that learning parameters fluctuate substantially over runs. Such fluctuations may limit the prospect of using single runs for individual diagnostics. Likewise, reliability of single-run estimates improved after several runs, highlighting the benefit of multi-run assessments to boost psychometric properties. Taken together, future use of innovative tools such as Influenca in large-scale, naturalistic assessments may provide a much more nuanced perspective on individual trajectories in decision-making and learning and their contribution to mental health.

Our app Influenca features several key innovations to study value-based decision-making and learning at a large scale as part of longitudinal studies in a naturalistic setting. Since the app was made available on the participants’ preferred devices and was completed outside of a controlled laboratory setting, a careful evaluation of the data quality is essential (Crump, McDonnell, & Gureckis, 2013; Germine et al., 2012; Gillan & Rutledge, 2021). To validate the online assessment, we show that participants quickly learn to do the task well and perform it in a moderately reliable manner after a few runs. Consequently, players win many more points than expected by chance in most of the runs 98% (score > .33; highest average win per trial when choices are simulated randomly) and wins increase over runs, indicating that participants played the task with increasing proficiency. In parallel with improvements in learning across runs, response times decreased, indicating that participants speed up their deliberation process with increasing proficiency as well. Beyond basic performance indices, we observed that participants showed changes in reinforcement learning parameters over runs. For example, the learning rate decreased on average over runs, indicating an integration of feedback about win probabilities over more consecutive decisions. In contrast, reward sensitivity increased over runs, reflecting greater exploitation of learned contingencies with increased task proficiency (Cohen, McClure, & Yu, 2007; Daw, O’doherty, Dayan, Seymour, & Dolan, 2006; Lee, Zhang, Munro, & Steyvers, 2011). Moreover, in line with previous findings (Fox & Poldrack, 2009; Kahneman & Tversky, 1980; Tom, Fox, Trepel, & Poldrack, 2007; Tversky & Kahneman, 1992), we observed that participants were more risk averse compared to a rational integration of relative value and probabilities. By design, the app does not promote a narrow range of learning rates and, instead, tries to capture individual differences in value-based decision-making over the course of the game. Such inter-individual variance is crucial for the effective use of tasks for individual prediction or classification (Hedge et al., 2018). To conclude, our newly developed app captures differences in inter-individual and intra-individual decision-making in a naturalistic setting and tracks increased proficiency and test-retest reliability of behavioral estimates over time.

Based on our extensive data of repeated runs of the task, we can provide a more refined insight into the test-retest reliability of value-based decision-making and learning. Across all runs, the reliability of the behavioral indices was poor to fair, indicating a limited trait-like characteristic of value-based decision-making (Neuser et al., 2020). However, the observed ICCs of the learning rate are in accordance with previous studies using lab-based assessments (Moutoussis et al., 2018; Pratt et al., 2020; Shiner et al., 2012) suggesting that this is unlikely due to the naturalistic setting. To see whether increased task proficiency would lead to improved reliability, we correlated estimates of single runs with the average of the held-out runs. Crucially, reliability of the learning rate increased up to Run 7, suggesting that initial variance in the parameter estimates is not necessarily as predictive of trait-like differences in learning as late variance in the presence of substantial task expertise. Therefore, multi-run assessments conducted across various states may provide a better approximation of generalizable inter-individual differences in value-based decision-making compared to the typical single-run assessments in the lab after limited practice on the task.

Despite its notable strengths, the study has several limitations that should be addressed in future research. First, in comparison to lab-based experimental setups, we cannot control the testing environment our participants are confronted with when they initiate the app. Still, we argue that the naturalistic setting of EMA has important advantages that can outweigh the limited control over standardized data collection, such as more data per person collected during representative circumstances they will face in their life outside the laboratory. Second, to illustrate the rationale of our app, we chose a reinforcement learning model suggested in the seminal work by Behrens and colleagues (Behrens et al., 2007). More advanced models could provide deeper insights into the decision-making processes and their progression over repeated runs. Potential advances include a hierarchical Bayesian framework (Mathys, Daunizeau, Friston, & Stephan, 2011), allowing to track the agent’s estimate of the current volatility in the environment, and asymmetric learning rates for wins and losses (Kravitz, Tye, & Kreitzer, 2012). Moreover, in our model fitting, we did not incorporate the nested data structure, which could be further exploited with Bayesian fitting methods to improve the estimation of trait-like characteristics of behavior (Boehm, Marsman, Matzke, & Wagenmakers, 2018). Third, apart from the rich possibilities provided by extensions in the computational models, future work may shed light on the reciprocal influence of momentary states (including metabolic states such as hunger) and the parameter estimates of value-based decision-making (Blain & Rutledge, 2020; Rutledge et al., 2017).

To summarize, online and smartphone-based assessment has gained traction as a scalable method for longitudinal studies in larger and more representative samples embedded in a naturalistic setting that may improve generalizability. Here, we provide a psychometric evaluation of our open source, cross-platform reinforcement learning task for future use in large-scale assessments of individual differences in value-based decision-making. We show that the gamified task captures inter-individual and intra-individual differences in decision-making and learning, which can be associated with fluctuations in state measures or potentially linked with interventional designs to provide more nuanced insight into behavioral changes. Based on our extensive longitudinal assessment of reinforcement learning over days, we provide detailed information on the test-retest reliability of the behavioral performance indices and model parameters, suggesting that multiple runs per participant might be necessary to provide sufficient diagnostic information at the individual level. To conclude, our reinforcement learning task can be used to precisely track the dynamics of value-based decision-making and learning providing a new avenue for future research to improve the individualized prediction of behavior as well as the classification of individuals for diagnostic or clinical applications.

## Code availability

The app was written in the open-source programming language HAXE (https://haxe.org/, Version 3.4.7). The source code for the Influenca framework is available via github (https://github.com/VTeckentrup/mind-mosaic). Compiled installers are available and will be maintained on our website www.neuromadlab.com/en/influenca-2.

## Acknowledgement

We thank Gizem Altan and Anastasia Illarionova for providing the graphics for the app, and Jennifer Them for support in the app programming. We thank Hannah Schütt for help in setting up the database. We thank Franziska Müller, Magdalena Ferstl, Salome Herwerth, Dana Wentz for help with data acquisition as well as Wiebke Ringels for support in additional analyses. The study was supported by the Else Kröner-Fresenius Stiftung, grant 2017_A67, and the Wikimedia foundation, open science scholarship granted to VT. VT & NBK received salary support from the University of Tübingen, Faculty of Medicine fortune grant #2453-0-0. MPN received additional salary support from the University of Tübingen, Faculty of Medicine ‘forschungsorientierte Gleichstellungsförderung’ 2605-0-0 awarded to NBK.

## Author contributions

NBK was responsible for the study concept and design. VT coded the task and provided technical support for users. MPN & VT collected data under supervision by NBK. NBK conceived the method. FK, MPN & AK processed the data. FK & AK performed the data analysis and VT, MPN, & NBK contributed to analyses. MPN, AK, FK & NBK wrote the manuscript. All authors contributed to the interpretation of findings, provided critical revision of the manuscript for important intellectual content and approved the final version for publication.

